## Supplementary figures and images for "Vertical transmission of maternal DNA through extracellular vesicles associates with altered embryo bioenergetics during the periconception period"

### Supp 1

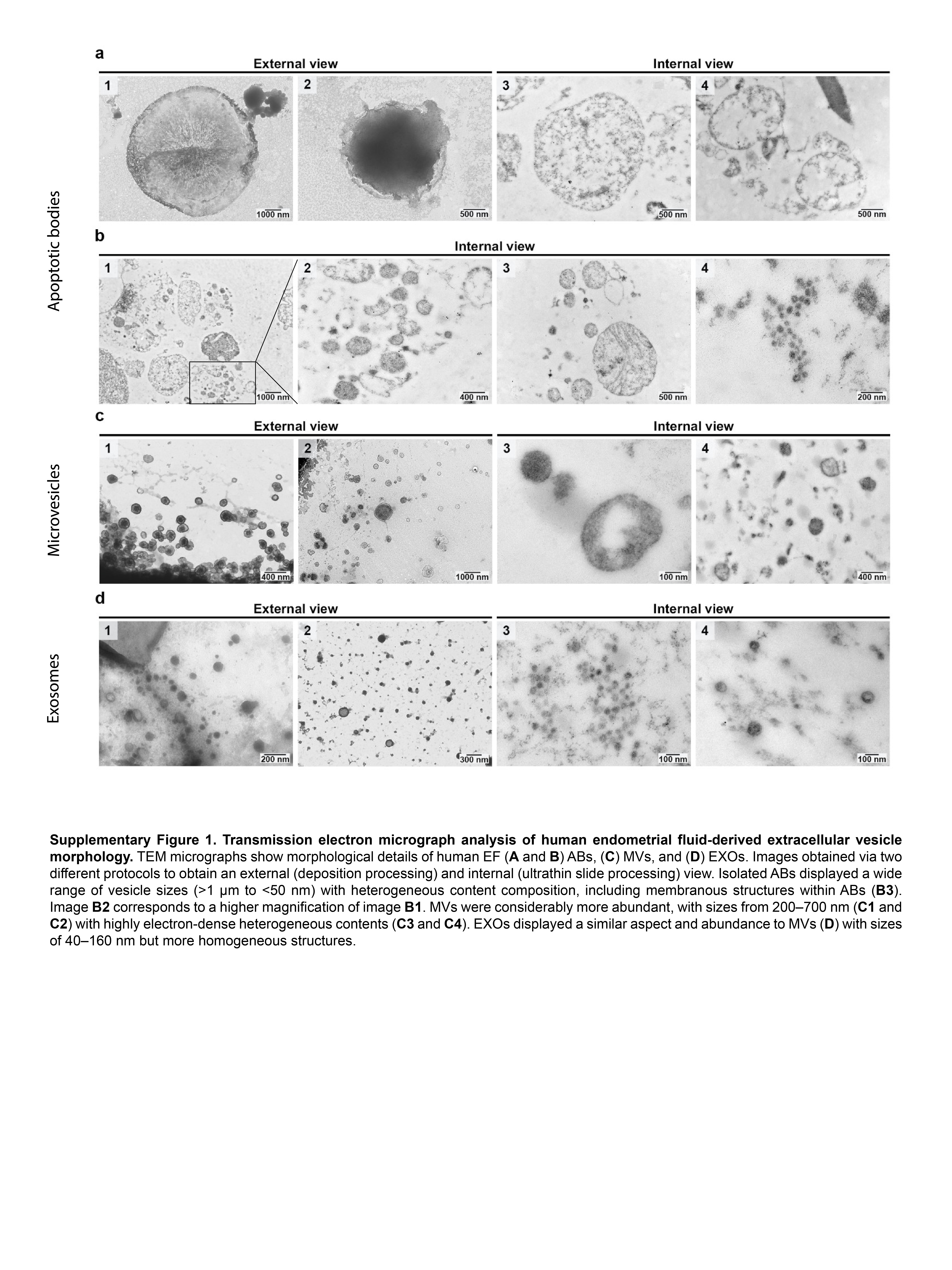

### Supp 2

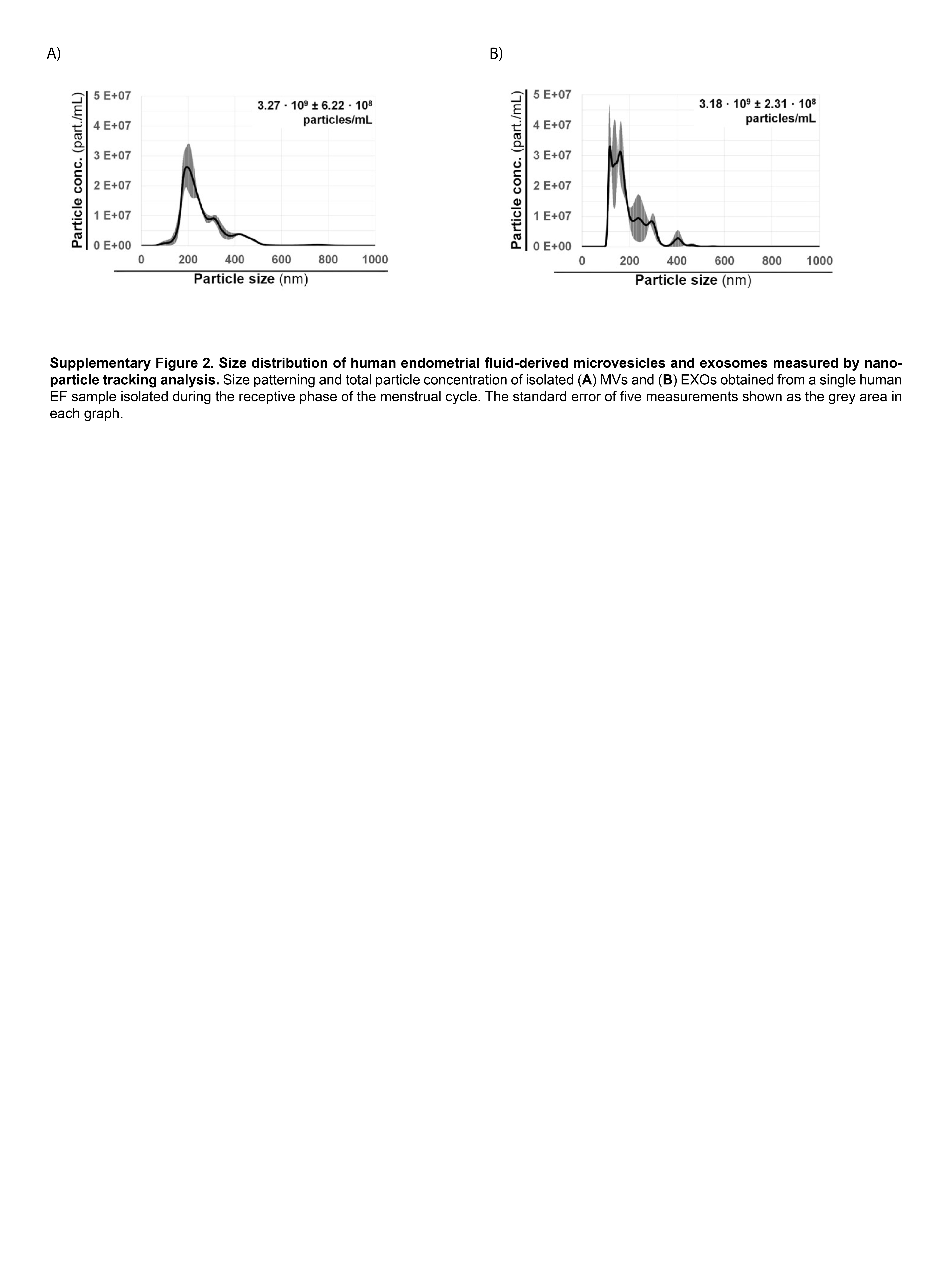

### Supp 3

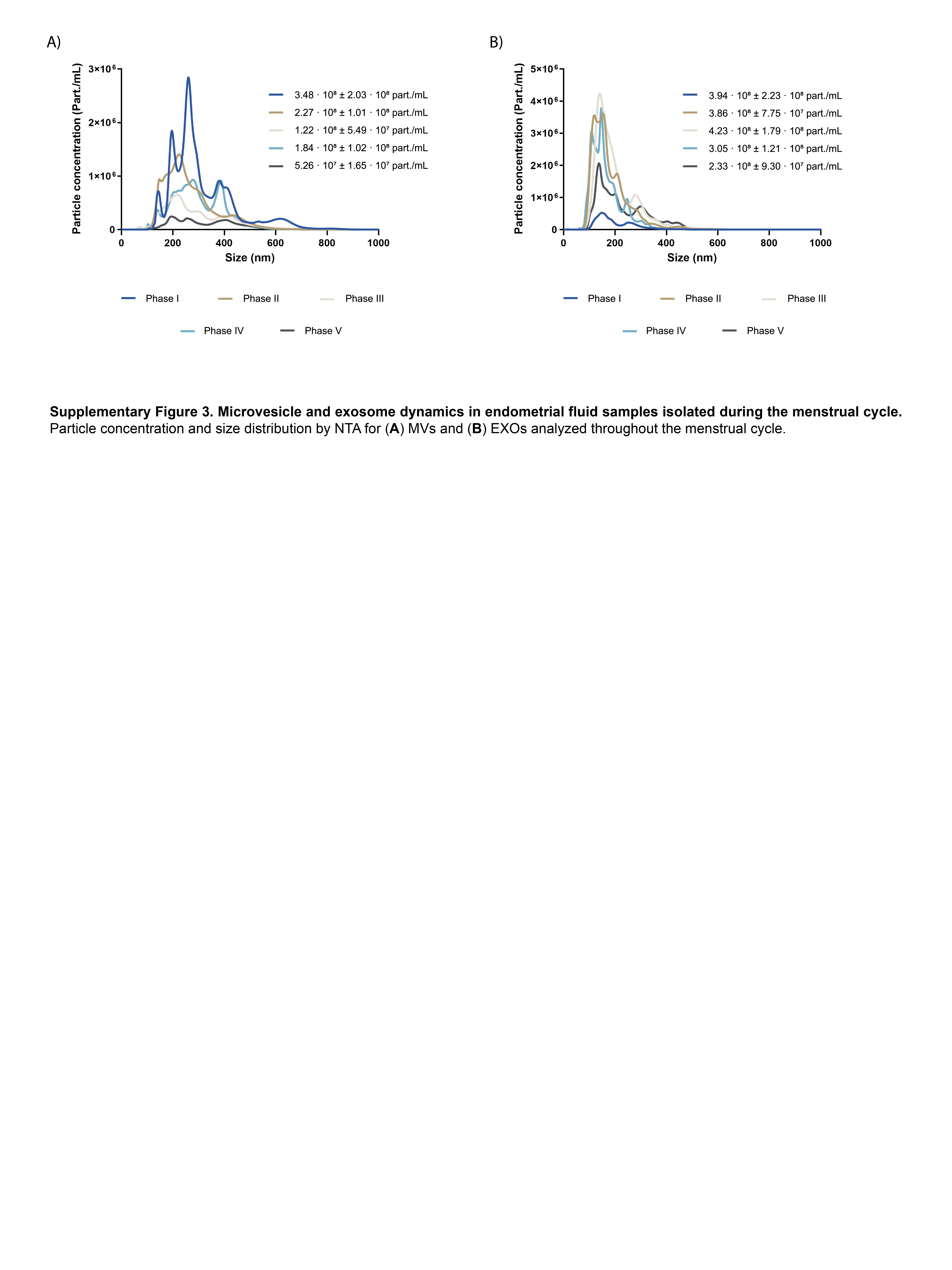

### Supp 4

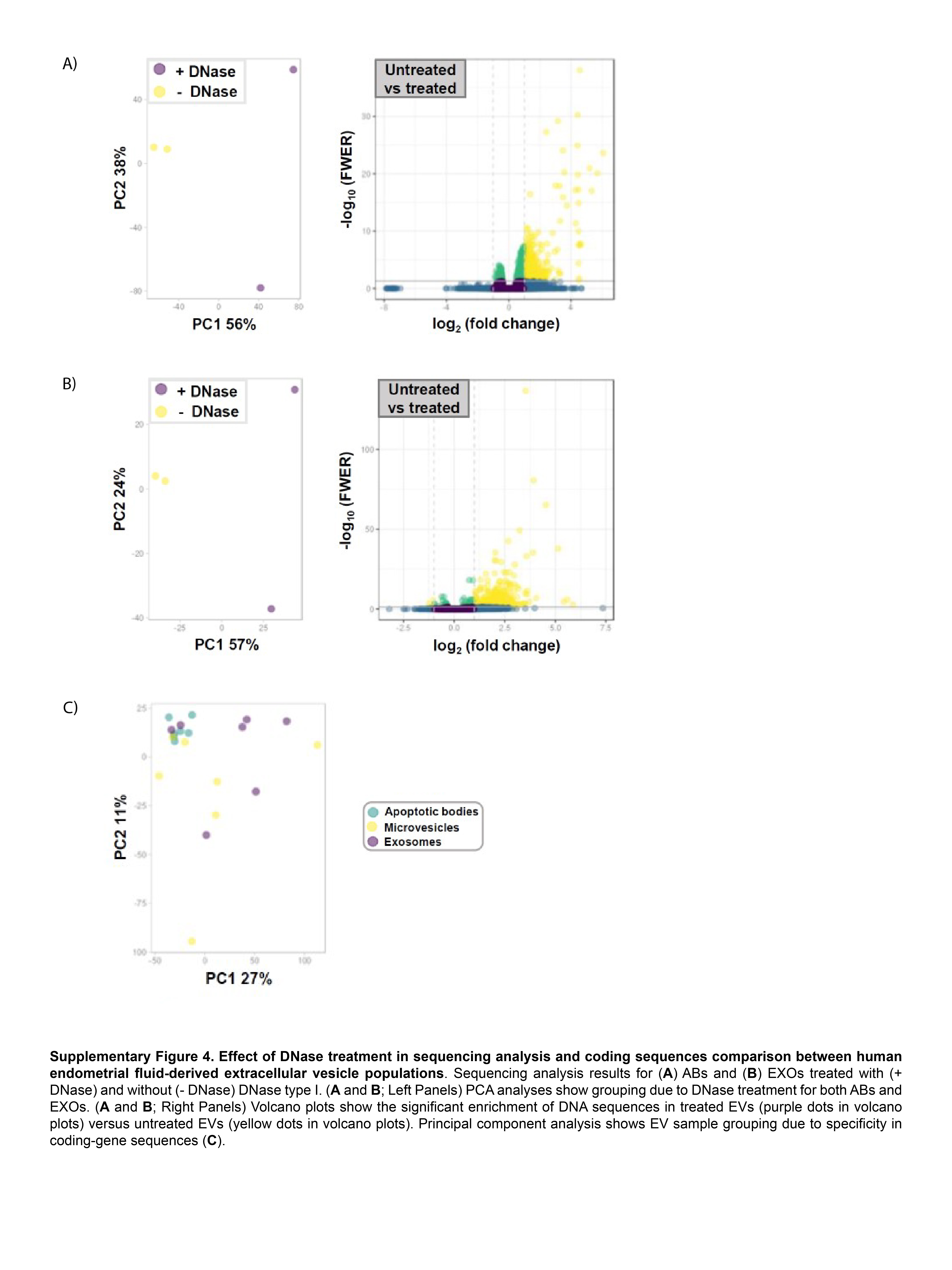

### Supp 5

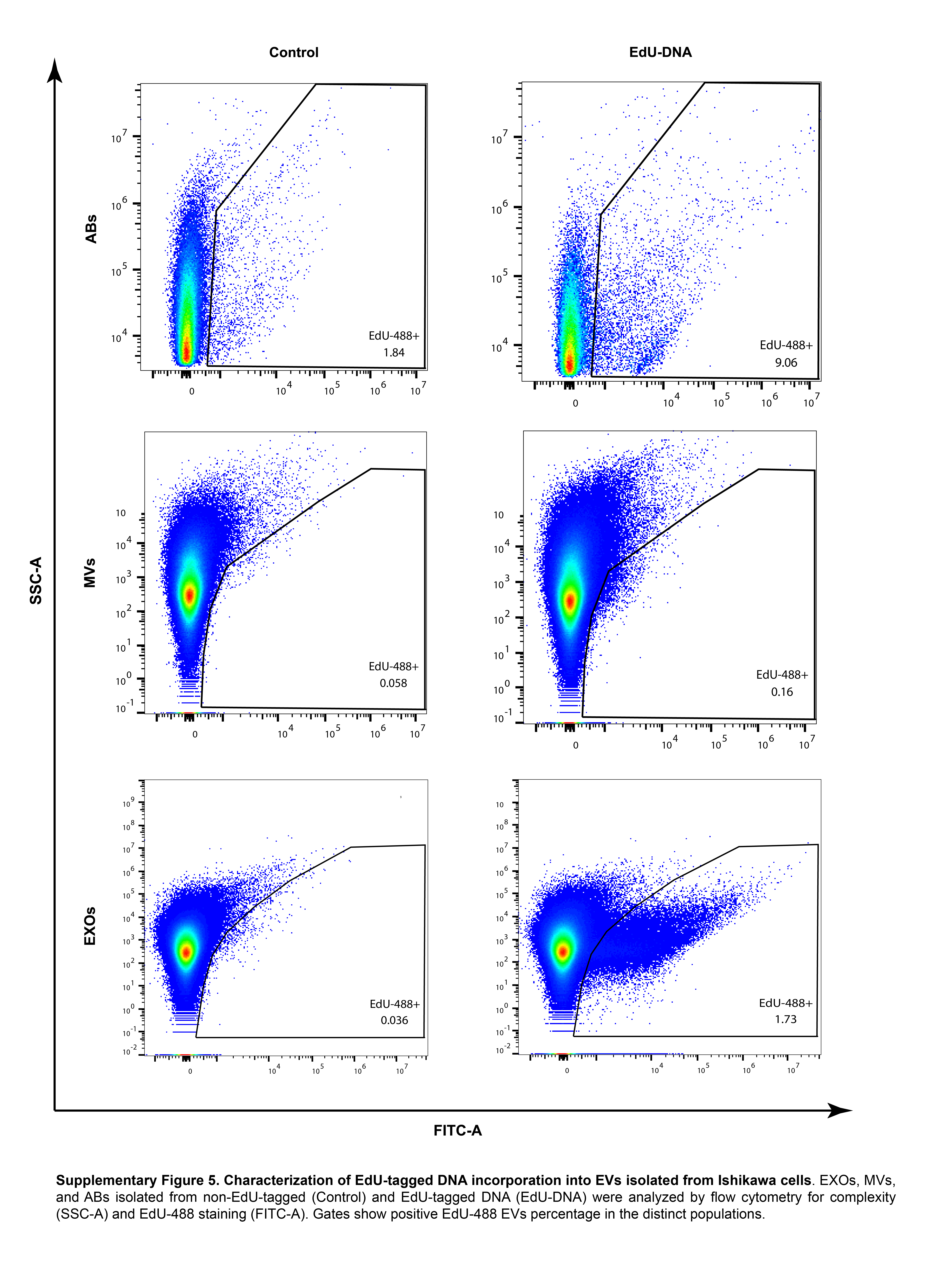

### Supp 6

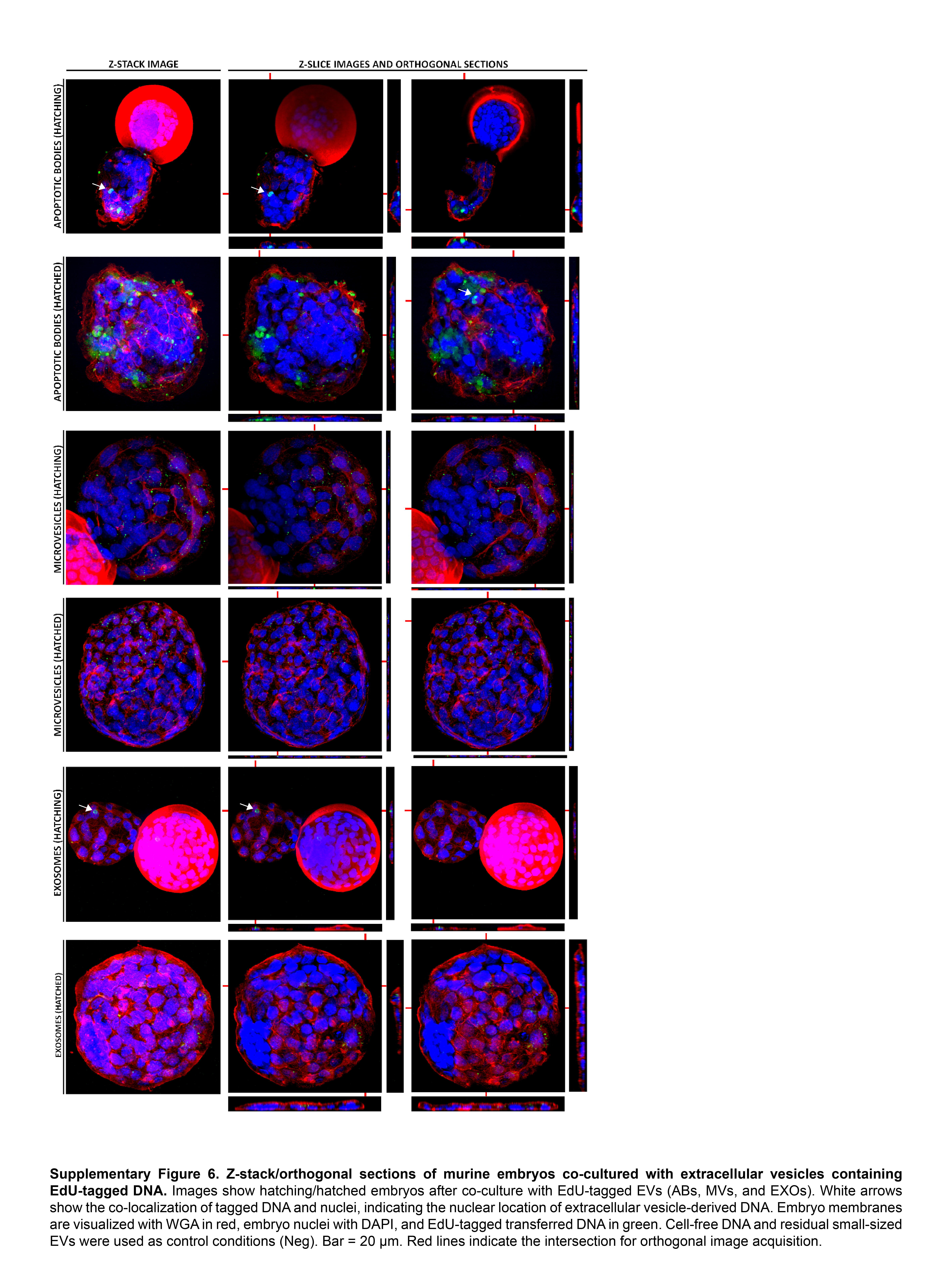

### Supp 7

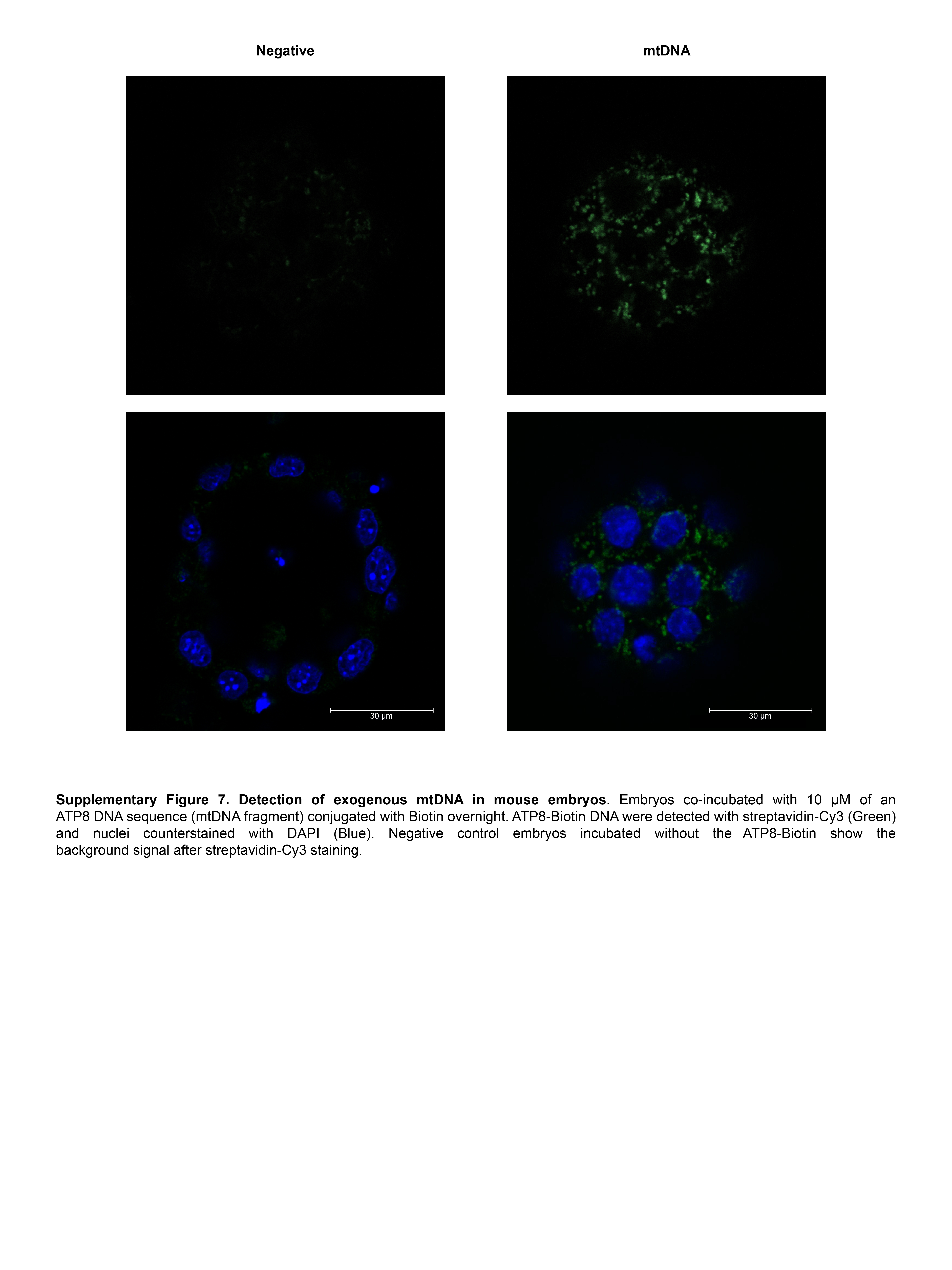
